# Validation of the Early Warning and Response System (EWARS) for dengue outbreaks: Evidence from the national vector control program in Mexico

**DOI:** 10.1101/2021.02.23.432448

**Authors:** David Benitez-Valladares, Axel Kroeger, Gustavo Sánchez Tejeda, Laith Hussain-Alkhateeb

**Author notes:** Corresponding author: Dr Laith Hussain-Alkhateeb;, c/o School of Public Health and Community Medicine, Institute of Medicine, Sahlgrenska Academy, University of Gothenburg, Box 414, 405 30 Göteborg, Sweden.

## Abstract

**Background:** During 2017, twenty health districts (locations) in Mexico implemented a dengue outbreak early warning and response system (EWARS) that uses epidemiological, meteorological and entomological variables (alarm indicators) to predict dengue outbreaks and triggers early response activities.

Eleven of these districts were analyzed as they presented reliable information. Nine districts presented outbreak alarms but without subsequent outbreaks (“non-outbreak districts”) and two presented after the alarms dengue outbreaks (“outbreak districts”). This study is concerned with i) if the alarms without outbreaks were false alarms or if the control services had established effective response activities averting an outbreak and ii) if vector control activities can mitigate or even avert dengue outbreaks.

**Methods:** Five components of dengue outbreak response (larval control, entomological studies with water container interventions, focal spraying, indoor residual spraying, space spraying) were quantitatively analyzed across two groups (”outbreak districts” and “non-outbreak districts”).

**Results:** The average coverage of vector control and responses were higher in non-outbreak districts and across all five components. In the “outbreak districts” the response activities started late and were of much lower intensity compared to “non-outbreak districts”. District vector control teams demonstrated diverse compliance with local guidlines for ‘initial’, ‘early’ and ‘late’ responses to outbreak alarms which could explain the different outcomes observed following the outbreak alarms.

**Conclusion:** findings from this study plausibly demonstrates important operational scenarios when succeeding or failing alarms signals generated by EWARS at national level. This study presents evidence warranting for further investigation into the effectiveness and cost-effectiveness of EWARS using gold-standard designs.

## Introduction

Dengue, a mosquito-borne viral disease, is currently one of the most important and fastest-spreading infectious diseases in the world, putting a huge burden on populations, health systems and economies in most tropical and sub-tropical countries (WHO, 2012). There are four viral serotypes circulating in Asia, the Americas and Africa with an estimated 3.6 billion people living in dengue endemic countries, and about 300 million infections occur annually (Bhatt et al., 2013). The growing global threat of dengue outbreaks in endemic and non-endemic regions of the world urges to focus at more effective outbreak management.

Routine passive surveillance systems for dengue and other infectious diseases are the backbone of epidemiological information in the vast majority of countries, allowing the analysis of the spatial and temporal distribution of clinically apparent dengue cases. These data can highlight "hot spots" and priority areas for interventions and serve as a trigger for local implementation of prevention and control activities. However, the underreporting of cases, especially of non-hospitalized dengue cases (in addition to asymptomatic or mild disease, non-users of the public health sector and others) is a serious problem. National surveillance systems have to use appropiate tools to trigger response actions for outbreaks and these must be sensitive to detect and potentially to predict outbreaks in a timely manner and, specific to prevent false outbreak alerts.

The early warning of outbreaks and the subsequent response is of vital importance to reduce, human suffering and economic losses both for families affected by the disease, as well as for health systems already weakened in most dengue endemic countries. An early warning system must typically be followed by a rapid outbreak response guided by standard operational procedures (SOPs).

The Special Programme for Research and Training in Tropical Diseases (WHO/TDR) initiated together with European research institutions, the national dengue control services and academia in endemic partner countries the development of a web-based Early Warning And Response System (EWARS) for dengue/ chikungunya/ Zika outbreaks (ref. Bowman 2016). Throughout multi-conuntry qualitative and quantitative assessments, the tool has revealed consistent outbreak predictions with instant interpretations including a subsequent action plan using the local vector control response protocol. The tool can be easily integrated into existing surveillance systems and based on previous reports the tool maintained high acceptability by users strengthening the communication and collaboration between central (national) level and district level as well as national and international partnerships (Hussain-Alkhateeb et al., 2018).

The centre for vector control at the Ministry of Health in Mexico (CENAPRECE) together with WHO TDR, and other endemic dengue countries (mainly Brazil and Malaysia) and national and international experts have been working since 2012 on the design and validation of the above mentioned early warning and response system for dengue outbreaks (EWARS, later also for chikungunya and Zika outbreaks; Cardenas et al. 2020) employing multiple epidemiological, meteorological and entomological indicators (WHO 2020 ; Bowman et al., 2016; Badurdeen et al., 2013).

In the context of EWS, interpreting essential terms such as “correct alarms” (alarm signals that were followed by an outbreak) and “false alarms” (alarm signals that were not followed by an outbreak) have significant local operational implications for a functioning surveillance system. Factors related to the timeliness and intensity of response activities, which tend to mitigate or avert the outbreak, can potentially be correlated with occasions of “no outbreak, no alarm” during the EWS process. This correlation, which primarily addresses the effectiveness of EWS in reducing unwanted disease outcomes, is rarely explored within the context of vector-borne diseases. To further ensure effective functions, EWS should be perceived as an information system designed to support decision making of national and local-level institutions, and able to improve coordination among relevant stakeholders, such as local epidemiologists, meteorologists, entomologists, the national and local management agencies that assess risk and develop response strategies, and the public communication channels used to disseminate warning information.

With this in view, the purpose of this study is to examine and compare outbreak response activities in districts with alarms followed by an outbreak (“outbreak districts”), and in districts which did not have an outbreak after the alarm (“non-outbreak districts”). We therefore hypothesized that delayed – or non-effective response activities measured by the timing (initial, early or late response) and the intensity of the response in terms of coverage or frequency of interventions – are associated with outbreak districts.

## Method

The prospective implementation of the EWARS tool in Mexico was carried out during 2015-17, but this analysis will focus on 2017 when the implementation was conducted with the highest degree of quality due to capacity building of surveillance staff in the study districts.

Of the 20 districts that participated in the implementation of EWARS (see map), only twelve presented alarms. Of those, only two had an outbreak after the alarm and nine did not during 2017, taking into account the operational definition of an outbreak (WHO and UNICEF, 2017).

The “outbreak districts” were Ciudad Apodaca and San Nicolás de los Garza (both in the State of Nuevo León). “Non-outbreak districts” were: Culiacán and Mazatlan (both in the State of Sinaloa), metropolitan area of Veracruz and Coatzacoalcos (both in the State of Veracruz), Apaztingan and Zamora (State of Michoacan), Cardenas (State of Tabasco), Iguala (State of Guerrero), Monterrey (State of Nuevo Leon).

In order to analyse the staged outbreak response (see table 1) (initial: *is declared when two consecutive alarm signals occur*; early: *is declared when three consecutive alarm signals occur*; and late response: *is declared when more than three consecutive outbreak weeks take place*) data were obtained on all dengue control activities during 2017 in the study districts from the online entomological surveillance platform of Mexico.

**Table 1.** Type of response and the corresponding dengue control activities

| Type of response | Initial response | Early response | Emergency or late response |
| --- | --- | --- | --- |
| **Dengue control activities** | Communicating risk to authorities, local dengue committees and outbreak management team at hospitals;<br>Updating the necessary background information such as cartography, demographics and inventory of facilities;<br>Focused vector control in areas considered historically high transmission. | Communicating increased outbreak risk to authorities, local dengue committees and outbreak management team at hospitals;<br>Convening local dengue or emergency operations committees;<br>Social mobilization (IEC, community participation, mass media partnerships);<br>Enhance communication channels with Public Health, Clinical, Education System., Media, national and international authorities; Vector control target the adult mosquito stages (focused around cases) with limited spatial fogging | Full application of the contingency plans. |

Essentially in regards to the time lag between the alarm and the response, due to the internal procedures of the independent Mexican epidemiological and entomological surveillance systems, it takes around one week to obtain the surveillance data and process the prediction model of the EWARS tool. Furthermore, dengue control activities are usually planned on weekly basis. Therefore, the time lag between an alarm and the response is usually two weeks, which was taken into account in this analysis.

The Mexican entomological surveillance platform divides response activities into five components: larval control, entomological studies, residual spraying around a probable case, indoor residual spraying and fogging (space spraying), and an indicator has been created for each response:

1. Larval control indicator: Number of houses covered per epidemiological week.
2. Indicator of entomological studies (which includes vector control activities): Number of entomological studies per epidemiological week.
3. Residual spray indicator around a probable case: percentage of probable cases covered with perifocal spray activities per epidemiological week.
4. Indoor residual spray indicator: number of houses covered per epidemiological week.
5. Fogging (space spraying) indicator: area covered (Km2) per epidemiological week

### Data management and statistical analysis

Due to inconsistencies in measuring the “*fogging (space spraying)*” during the data collection process, we only analyzed and discussed the other four response indicators quantitatively with concluding remarks about the realities of space spraying (fogging). The coverage of vector control activities (i.e. % of houses covered) at each district was calculated for each one of the four vector control activities before and after the alarm signal period (week). The coverage was calculated by dividing the total number of worked households by the total number of households in the corresponding district for the *larval control* and the *indoor residual spraying control*. The percentage of the number of *entomological studies* was estimated in relation to the total number of households of the corresponding district (see comments on coverage above). For the *spraying around new cases* (“perifocal spraying”), this indicator was computed by dividing the number of houses with (probable) dengue cases sprayed in relation to the total number of houses with probable cases reported in the corresponding district.

Segmented time-series analysis was used to examine the trend of coverage of vector control activities at baseline (the period prior to the EWS alarm signal), at the time of the alarm signal (this measures the immediate change in vector response between the baseline period and one week after the alarm signal) and, at the remaining period (weeks) following the alarm signal for each district. These three different time-periods trends were presented as “rates of change in percentage or per 1000 population” together with their p-values at 5% significant cut off. Descriptive statistics, in numbers and graphs, were also produced and presented before- and after- the alarm signal week, generated from the EWARD, for the corresponding district and indicator.

We developed two graphical representations of pre- and post-alarm activities, in the example larval control. Fig 2a shows the weekly coverage rates achieved following the alarm signal (green line). Fig 2b shows the trends of activities before and after the alarm signal became positive based on the segmented time series analysis (see below). However, the tables resulting from the segmented time series analysis gave a more complete picture, so we decided to use these for the data analysis and presentation.

**Figure 1.**
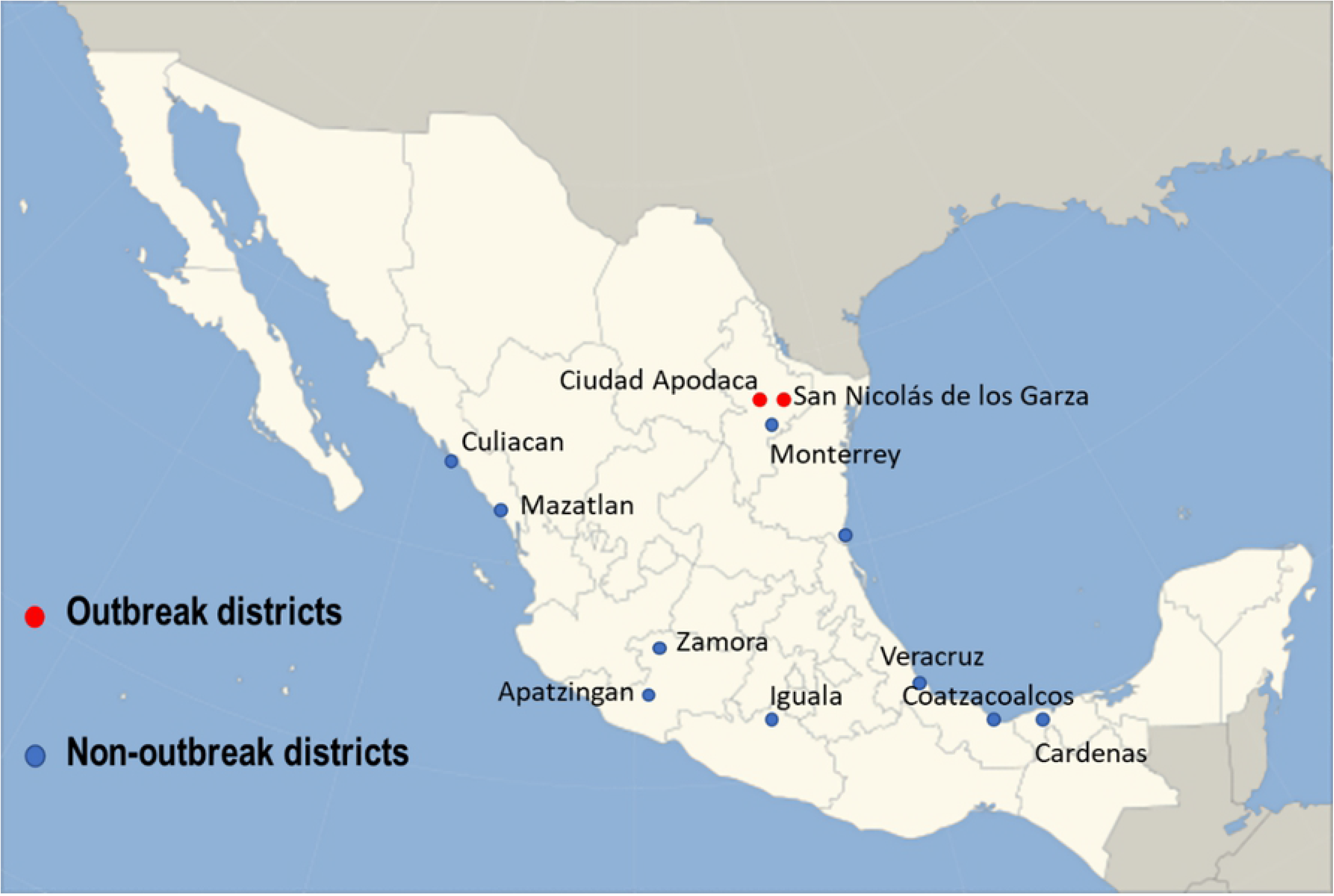
Map of Mexico illustrating the Intervention and non-intervention districts

**Figure 2a:**
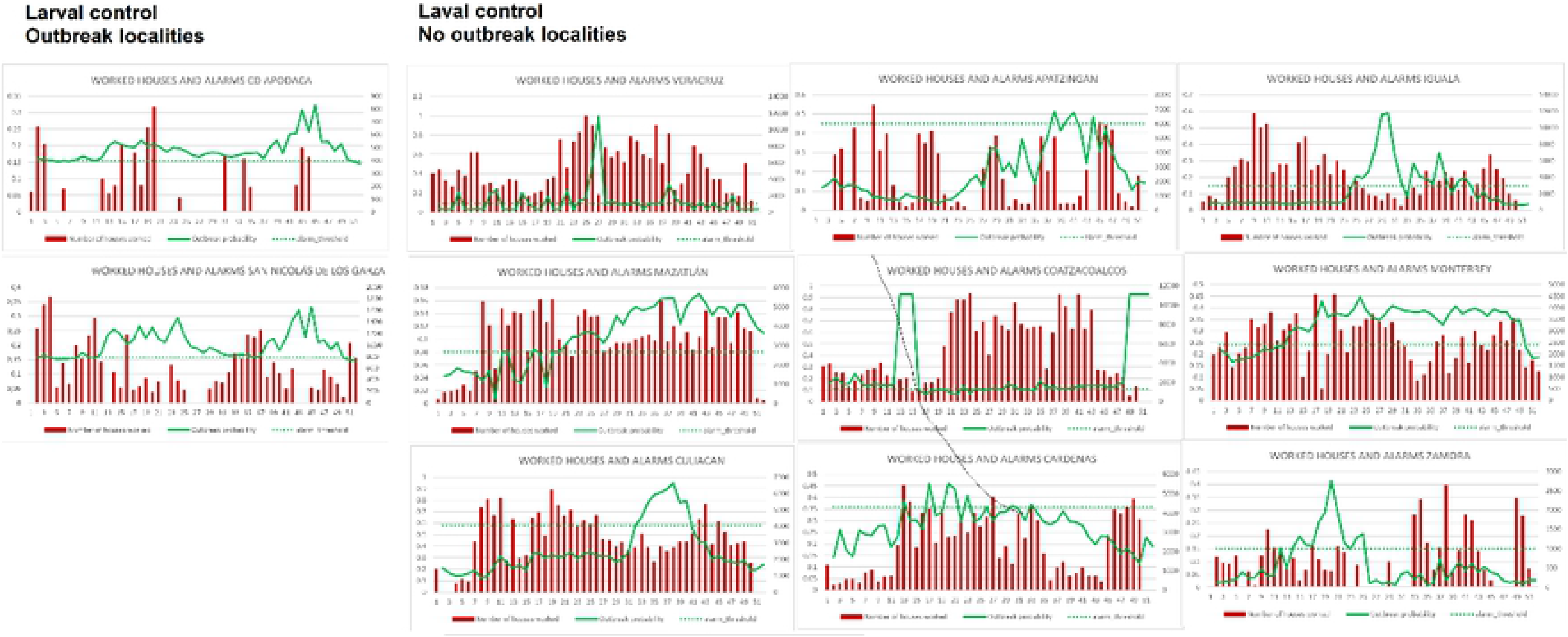
Weekly number of houses covered with larval control before and after outbreak alarm

**Figure 2b:**
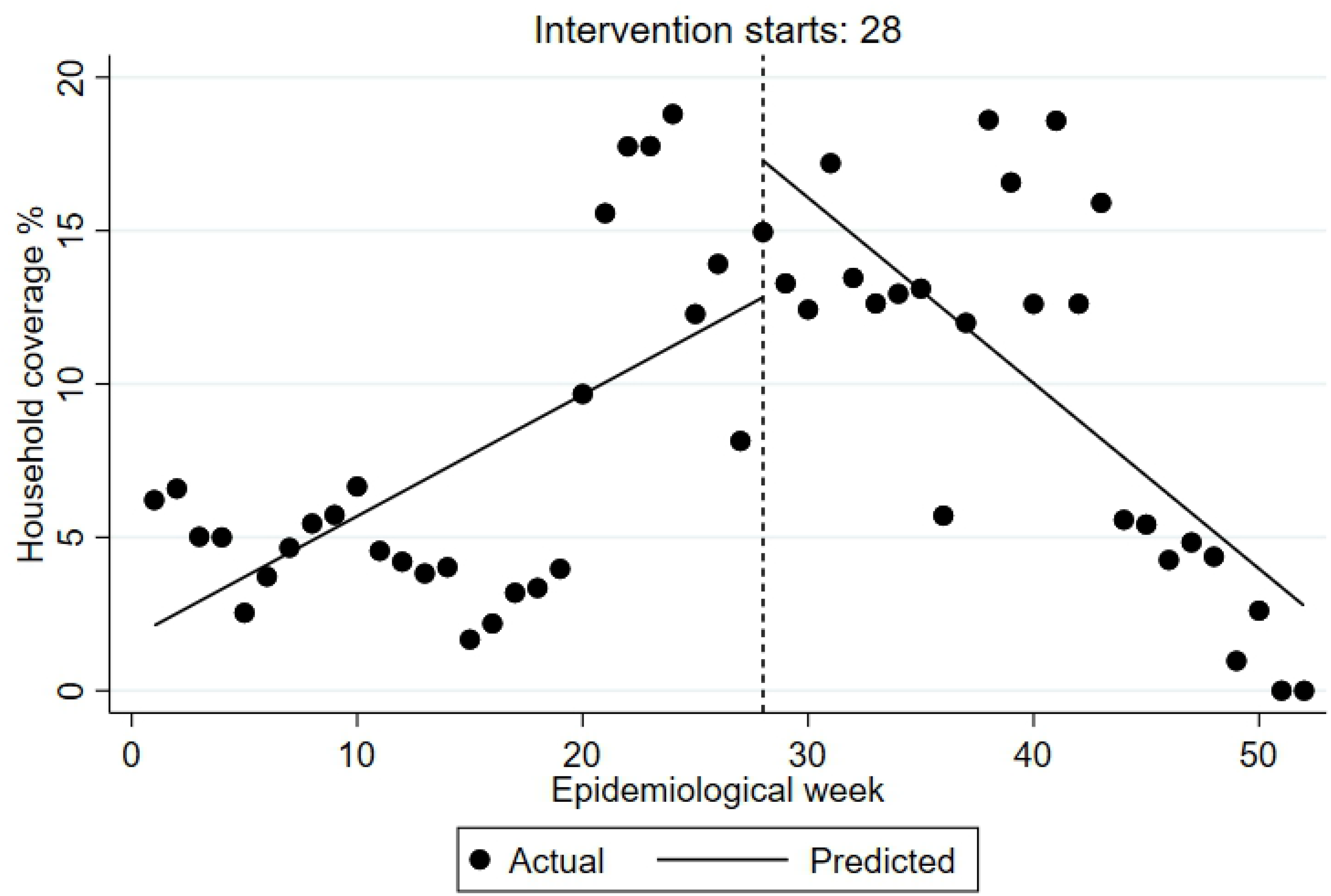
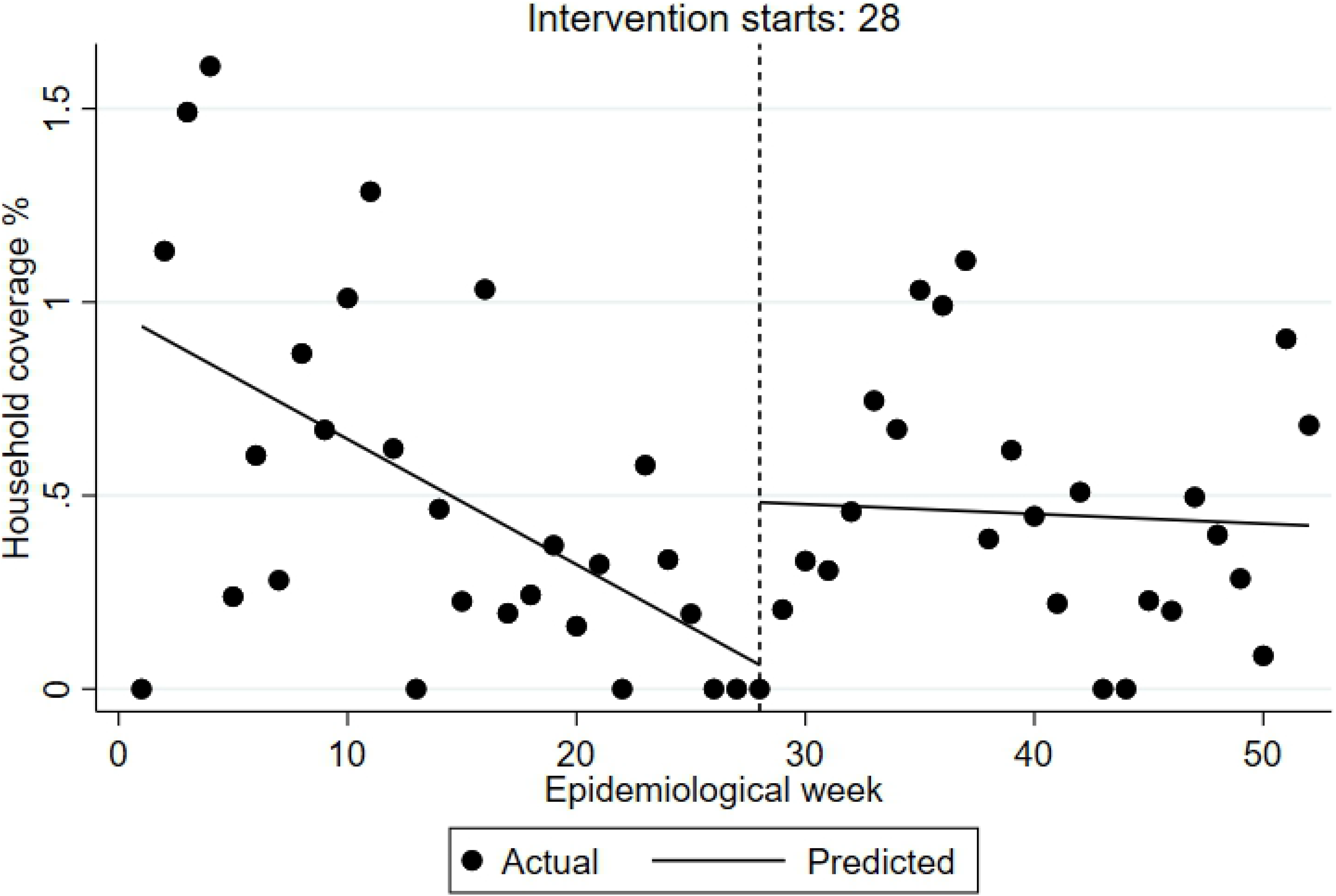
Weekly number of houses covered with larval control before and after outbreak alarm resulting from the segmented time series analysis. Two examples, one of outbreak and another of non-outbreak districts are presented. Coatzacoalcos - a non outbreak district San Nicolas - an outbreak district

#### Ethics approval

Ethical approval was obtained by the Pan American Health Organization Ethics Review Committee (PAHO ERC 2011-12-021)

## Results

Figure 3 demonstrates the outbreak prediction scenario of the corresponding district. The endemic channel for each epidemiological week during 2017 with the upper line being the outbreak threshold is illustrated together with the alarm threshold and the estimated outbreak probability by the EWS using climatic, epidemiological and entomological indicators. Alarm signals are triggered once the outbreak probability (“alarm line”) exceeds the alarm threshold line indicating a probably outbreak. The weekly dengue incidence rates are also shown to inform of the propospective distribution of cases in relation to historical incidences (endemic channel). In “outbreak districts”, the alarm signal is followed by an outbreak, whereas in the “non-outbreak districts”, the alarm signal is not followed by an outbreak.

**Figure 3.**
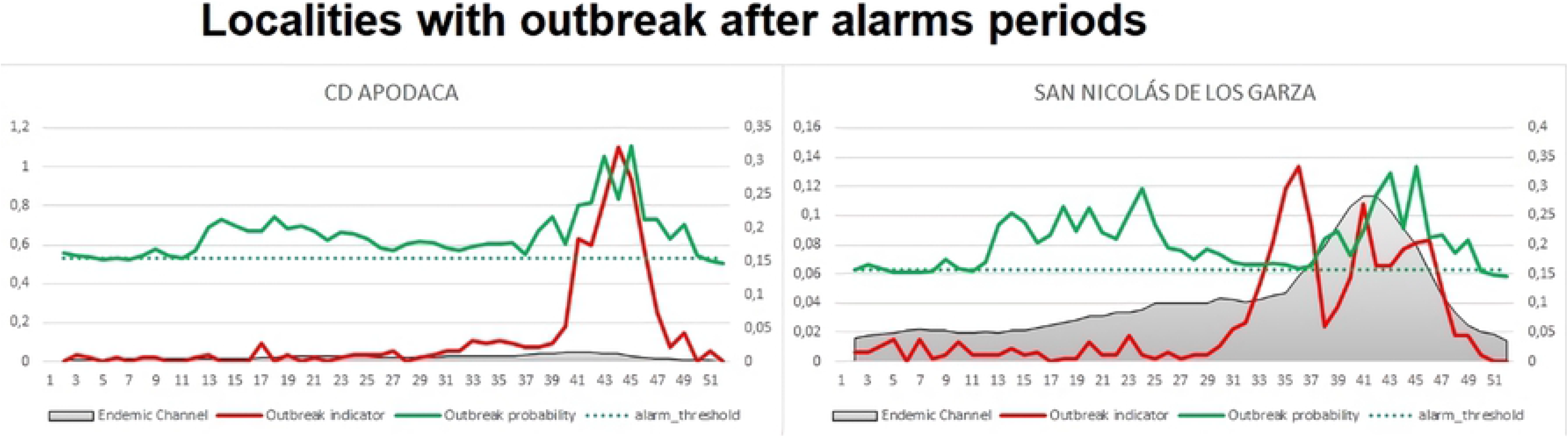
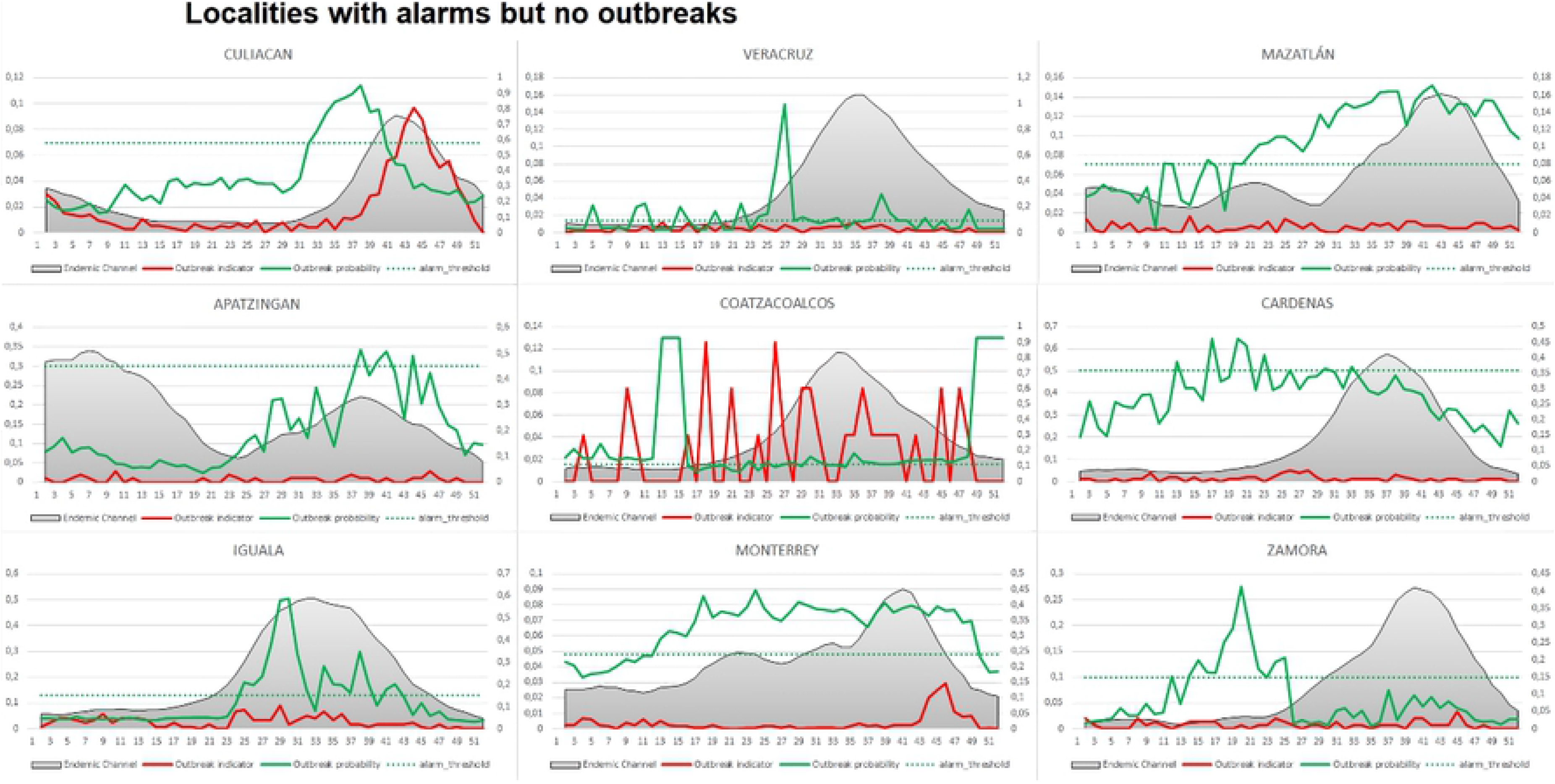
the outbreak predictions as generated by the EWARS for all nine districts. Footnote: The endemic channel is presented by the shaded area in grey which includes + z* SD of the moving average; the alarm threshold is represented by the dotted horizontal line; the outbreak probability (“alarm line”) estimated by the EWS is represented by the green line; the weekly dengue incidence rates is represented by the red line.

### Vector control indicators in outbreak and non-outbreak districts

#### Larval control indicator

Table 2 presents the findings from the segmented time-series analysis and the percentage of household coverage at district-level for the larval control (see comments on coverage estimates in the introduction).

**Table 2.**
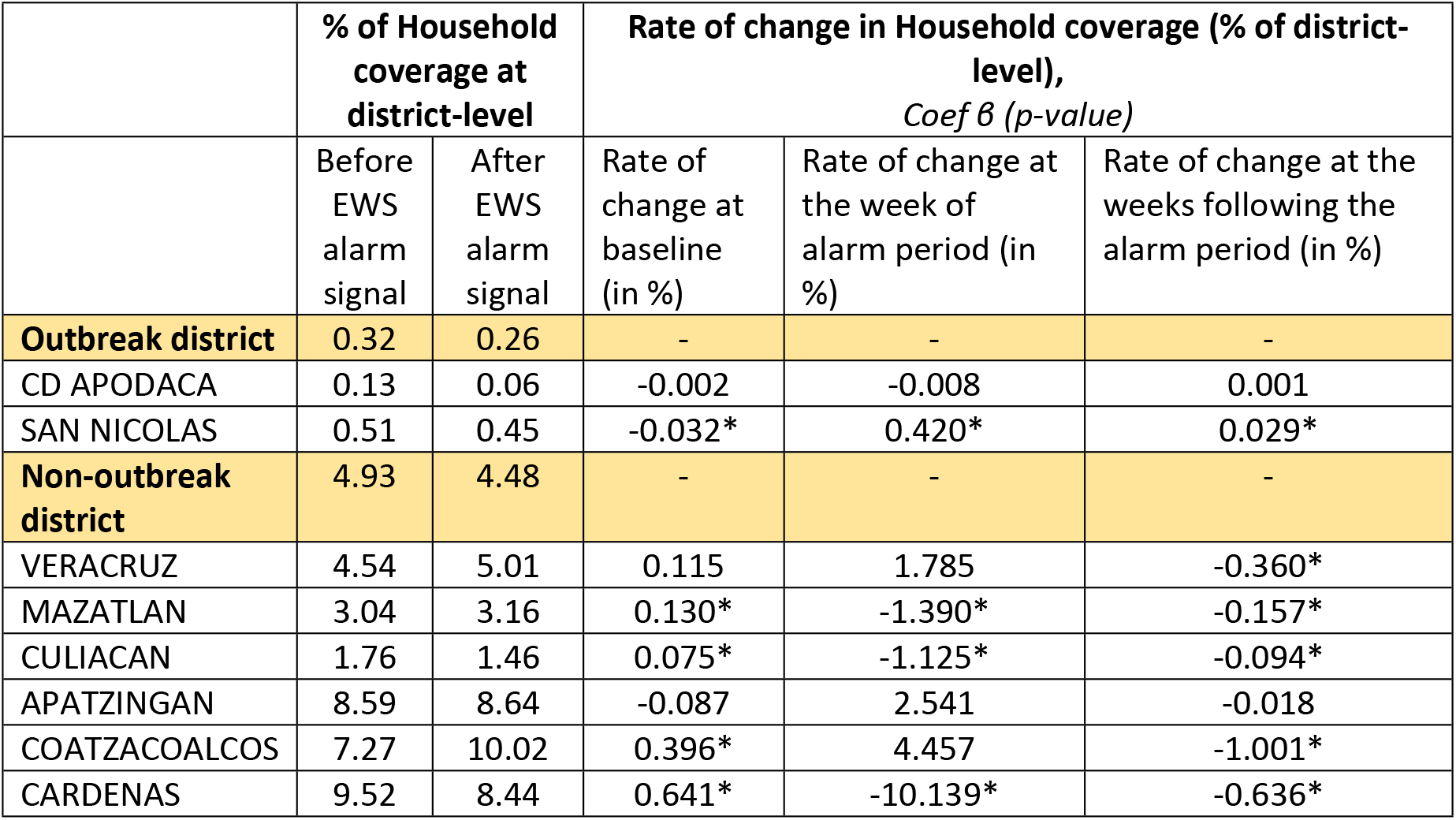
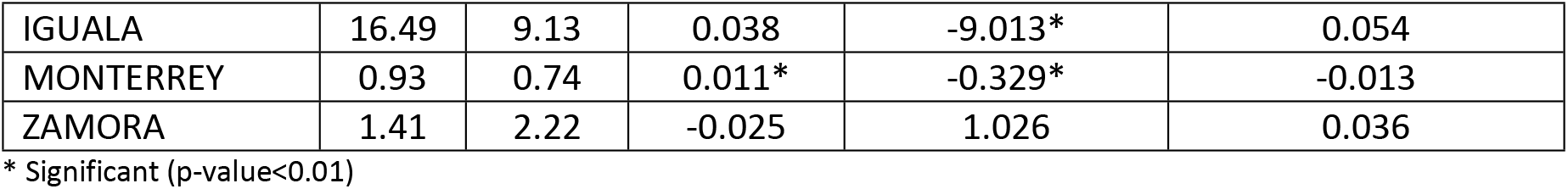
**Intensity of larval control** before and after the alarm signal across 2 outbreak-districts and 9 non-outbreak districts with the rate of change (from before to after alarm period) for vector control activity changes.

The coverage at household level was clearly much higher in non-outbreak districts (compared to outbreak districts) but it also varied considerably between districts with the highest coverage amongest the non-outbreak districts. From the segmented analysis, it is apparent that the rate of larval control was consistently declining among the outbreak districts and only slightly increased following the alarm signal generated by the EWS. The non-outbreak district group, however, maintained a routine increase of larval control rate at baseline (i.e. before any alarm occurred) with the exception of two districts out of nine (Apatzingan and Zamora). In contrast, the outbreak district group attempted to slightly increase the rate of larval control only after the alarm signal turned positive but particularly after the case numbers increased (see table 1). Most of the non-outbreak districts tended not to intensify or even reduce their larval control activities putting more emphasis on other outbreak response activities.

#### Entomological studies

In Mexico, vector control personnel perform entomological studies on a sample basis to control the quality of larval control activities and at the same time enhance larval control.

According to the segmented time series analysis the intensity and distribution of entomological studies followed a fairly similar pattern as observed for the larval control indicators. The coverage in general was lower as the entomological studies are being done on a sample basis. However, the non-outbreak districts maintained a higher percentage of entomological studies compared to the outbreak districts in general. Unlike the outbreak districts, the non-outbreak group showed an increased rate of activities at baseline, i.e. prior to any alarm signal, with exception of two districts. The rate of entomological studies varied across the groups at the post alarm period but remained at a low rate with the exception of the non-outbreak districts Apartzingan, Coatzacoalcos and Zamora and one outbreak-district (SAN NICOLAS) which showed an increasing trend after the alarm signal but starting late (see above); see table 3.

**Table 3.**
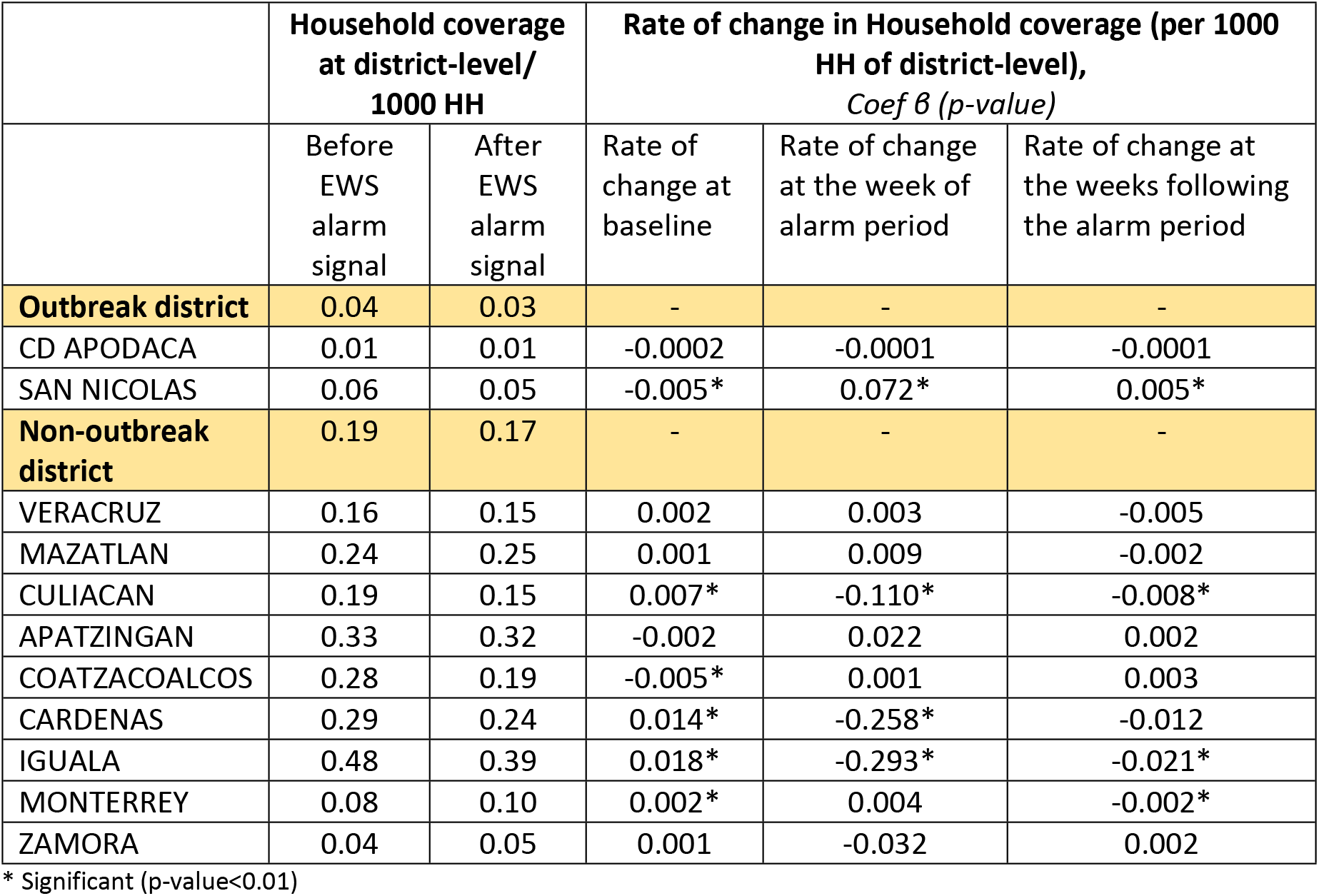
**Intensity of Entomological studies** before and after the alarm signal across 2 outbreak-districts and 9 non-outbreak districts with the rate of change (from before to after alarm period) for vector control activity changes.

|  | Household coverage at district-level/<br>1000 HH |  | Rate of change in Household coverage (per 1000 HH of district-level),<br>*Coef $\beta$ (p-value)* |  |  |
| --- | --- | --- | --- | --- | --- |
|  | Before EWS alarm signal | After EWS alarm signal | Rate of change at baseline | Rate of change at the week of alarm period | Rate of change at the weeks following the alarm period |
| **Outbreak district** | 0.04 | 0.03 | - | - | - |
| CD APODACA | 0.01 | 0.01 | -0.0002 | -0.0001 | -0.0001 |
| SAN NICOLAS | 0.06 | 0.05 | -0.005* | 0.072* | 0.005* |
| **Non-outbreak district** | 0.19 | 0.17 | - | - | - |
| VERACRUZ | 0.16 | 0.15 | 0.002 | 0.003 | -0.005 |
| MAZATLAN | 0.24 | 0.25 | 0.001 | 0.009 | -0.002 |
| CULIACAN | 0.19 | 0.15 | 0.007* | -0.110* | -0.008* |
| APATZINGAN | 0.33 | 0.32 | -0.002 | 0.022 | 0.002 |
| COATZACOALCOS | 0.28 | 0.19 | -0.005* | 0.001 | 0.003 |
| CARDENAS | 0.29 | 0.24 | 0.014* | -0.258* | -0.012 |
| IGUALA | 0.48 | 0.39 | 0.018* | -0.293* | -0.021* |
| MONTERREY | 0.08 | 0.10 | 0.002* | 0.004 | -0.002* |
| ZAMORA | 0.04 | 0.05 | 0.001 | -0.032 | 0.002 |

\* Significant (p-value<0.01)

Regarding the timeliness of the response activity “entomological studies” (figure 4), there was mainly a late (or emergency) response in both outbreak districts while in six out of nine non-outbreak districts there was an increase of entomological studies as initial response, followed by more extensive routine activities. In the other districts, however, there was an intial response followed by routine entomological studies.

**Figure 4.**
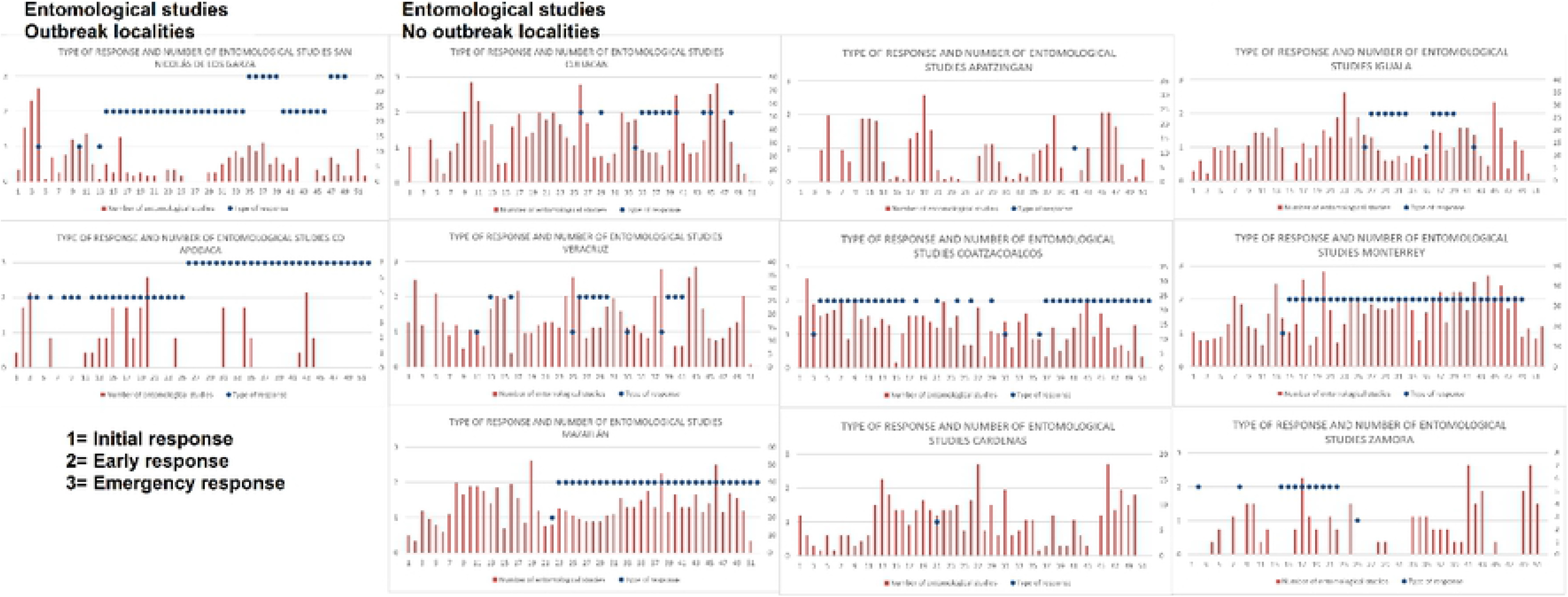
Timelines of the responses. Illustration of initial, early and late (emergency) responses as practiced in all nine districts based on the prediction generated from the EWARS.

#### Spraying around a case (Perifocal spraying)

The same pattern as above can be observed regarding perifocal spraying around a suspected dengue case. There was no or very little activity in outbreak districts but a quite frequent routine perifocal spraying in the non-outbreak districts, often enhanced after an outbreak alarm (Table 4).

**Table 4.**
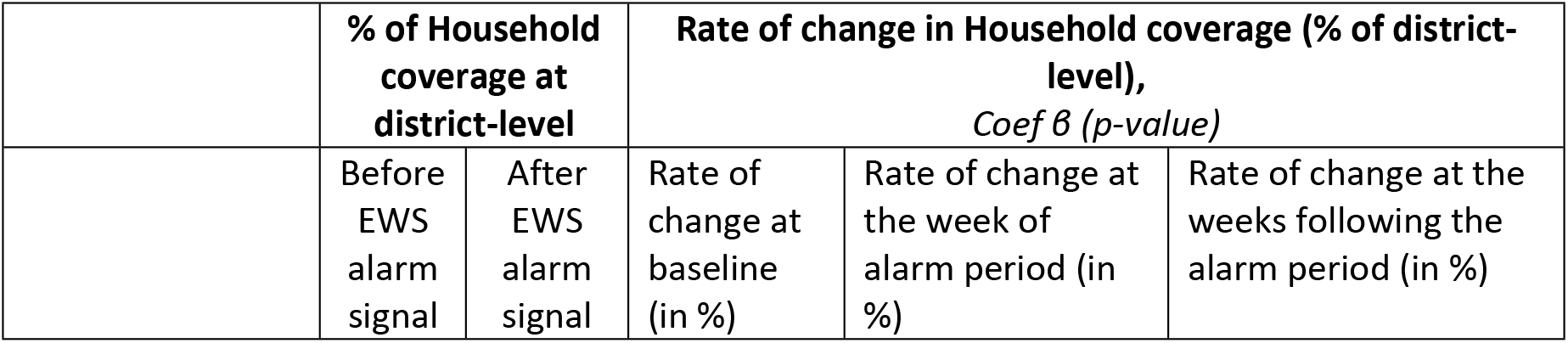
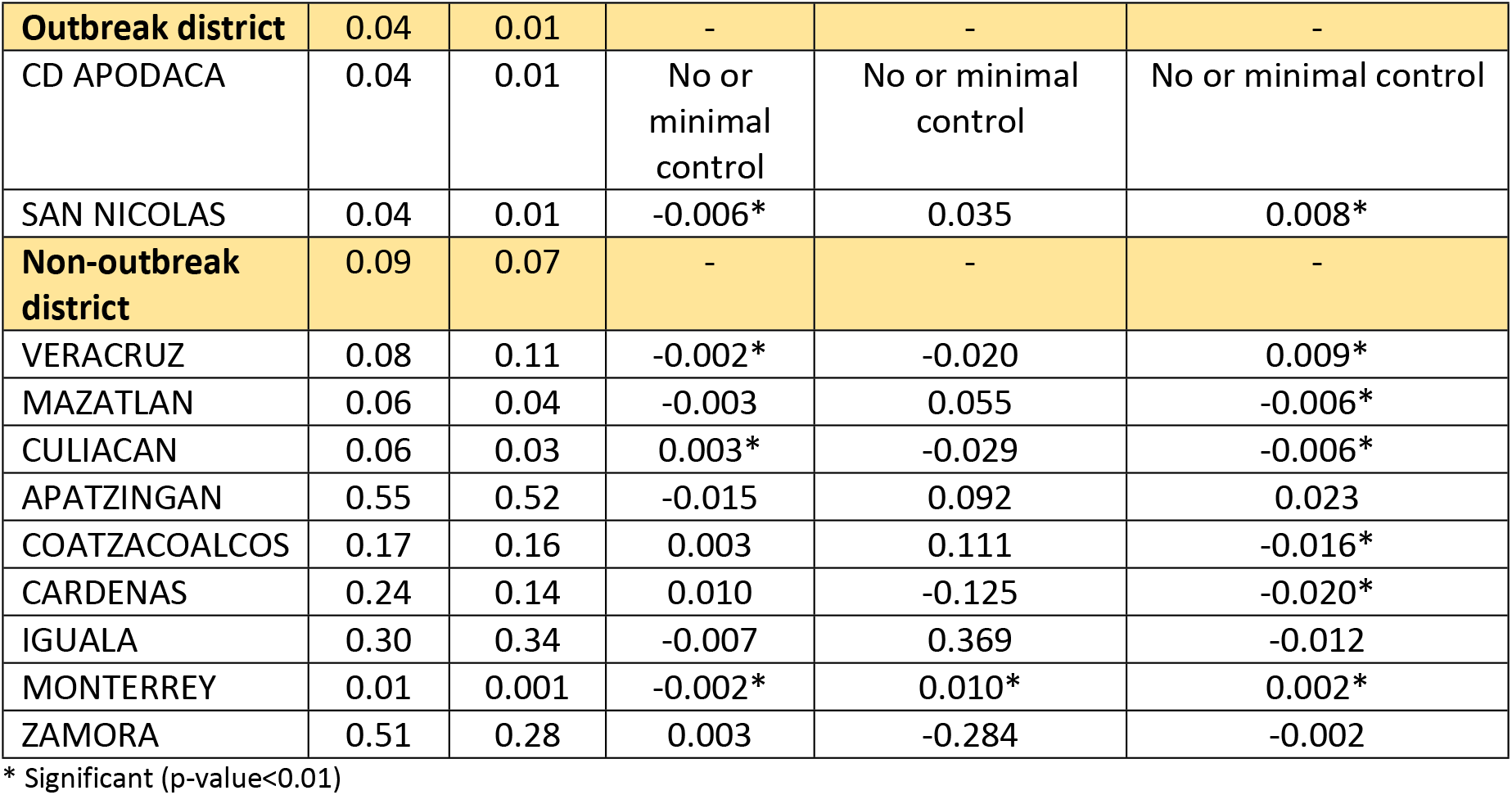
**Intensity of Spraying around cases (perofocal spraying)** before and after the alarm signal across 2 outbreak-districts and 9 non-outbreak districts with the rate of change (from before to after alarm period) for vector control activity changes.

Regarding the timeliness of the response (figure 3), no response could be observed in outbreak districts. In contrast, in six of the nine non-outbreak districts increased perifocal spraying as initial response was observed followed by continued early response activities. In the other three non-outbreak districts initial response was continued with routine activities. This pattern was confirmed by findings from the segmented time-series analysis (table 4). The coverage of spraying around cases was low in the outbreak districts and this continued showing no or minimal rates of change before and after the alarm signal, with the exception of one outbreak-district (San Nicolas). Spraying around cases in non-outbreak districts varied considerably from one district to another, and this was reflected in the trend at baseline, at the time of the alarm signal and during the period after the alarm, where most districts chose to retain low activities.

#### Indoor residual spraying (IRS)

This indicator also shows that response activities – indoor residual spraying, Table 5 – in outbreak districts were only carried out once the outbreak had begun. In most non-outbreak districts – at baseline before the alarm signal predicted an outbreak – there were already considerable increasing indoor spraying activities going on with the exception of two districts. After the alarm, there was an increase in response activities with indoor spraying..

**Table 5.** **Intensity of indoor spraying control** before and after the alarm signal across two outbreak-districts and nine non-outbreak districts with the rate of change (from before to after alarm period) for vector control activity changes.

| | % of Household coverage at district-level | | Rate of change in Household coverage (% of district-level),<br>Coef $\beta$ (p-value) | | |
| --- | --- | --- | --- | --- | --- |
|  | Before EWS alarm signal | After EWS alarm signal | Rate of change at baseline (in %) | Rate of change at the week of alarm period (in %) | Rate of change at the weeks following the alarm period (in %) |
| **Outbreak district** | 0.01 | 0.06 | - | - | - |
| CD APODACA | 0.01 | 0.02 | 0.0001 | 0.012 | 0.0001 |
| SAN NICOLAS | 0.02 | 0.11 | 0.0001 | 0.091* | -0.001 |
| **Non-outbreak district** | 0.29 | 0.59 | - | - | - |
| VERACRUZ- | 0.62 | 0.47 | 0.023 | -0.086 | -0.056* |
| MAZATLAN- | 0.02 | 0.03 | 0.002* | 0.002 | -0.003* |
| CULIACAN | 0.02 | 0.05 | 0.001* | 0.042* | -0.003* |
| APATZINGAN | 1.25 | 3.79 | 0.062 | 2.256* | -0.109 |
| COATZACOALCOS | 0.01 | 0.01 | 0.0001 | 0.001 | -0.001 |
| CARDENAS | 0.06 | 0.01 | -0.009* | 0.086* | 0.009* |
| IGUALA | 0.77 | 1.22 | 0.105* | -1.015* | -0.104* |
| MONTERREY | 0.02 | 0.08 | -0.002 | 0.030 | 0.003 |
| ZAMORA | 0.44 | 0.66 | 0.028* | 0.002 | -0.043* |

\* Significant (p-value<0.01)

Regarding timeliness of the response (figure 3), in the outbreak districts the increase of indoor spraying occurred long after the alarm and only when the case numbers went up. In five of nine non-outbreak districts initial response activities were started after the alarm signal, followed by early response activities. This is also illustrated in Table 5 revealing a general low coverage of indoor spraying activities (as this is a very resource intensive activity) but being significantly higher in four out of nine non-outbreak districts. Unlike the case in the other three response activities, most of the districts in both the outbreak and non-outbreak groups demonstrated increasing trends of indoor spraying from baseline to the time of the alarm signal.

#### Space spraying (fogging)

Despite some inconsistencies in measuring the “fogging (space spraying)” during the data collection process that prevent quantitative analysis, a qualitative analysis is outlined instead. For the space spraying, the worked area indicator (Km2) has been used per epidemiological week. This indicator shows a somewhat different pattern from the rest of the components. There is an increase in activities in the alarm periods in the locations with an outbreak, while, on the other hand, it could be said that this increase is not so large in those locations that did not show an outbreak (see figure 5). The case of Culiacan, a non-outbreak district, is emblematic since it presents a decrease in the activity of space fogging just after the alarm period. In the other localities that did not present an outbreak, this decrease does not exist as such, although the increases in the activity referred to in this component of the dengue response are similar to the increases experienced in the localities that did suffer an outbreak during 2017.

**Figure 5.**
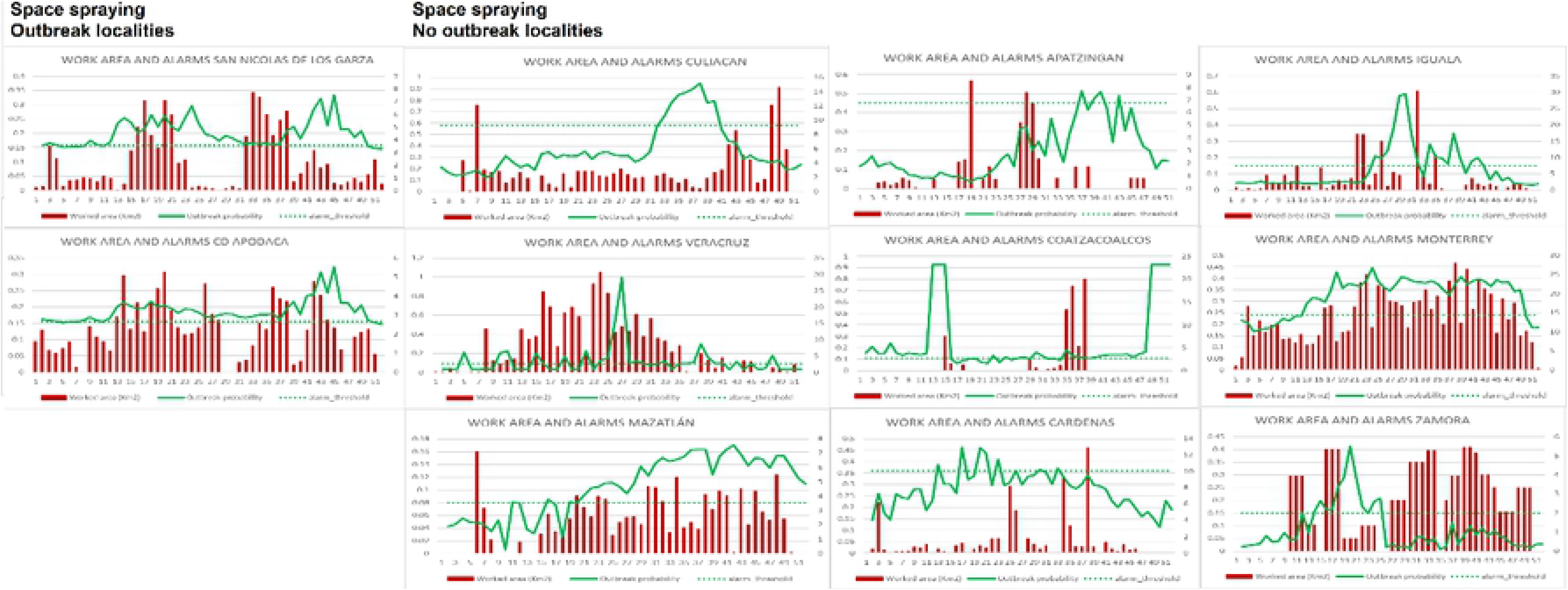
Space spraying weekly activities across outbreak and non-outbreak districts in 2017

Although there is an increase in fogging activities in San Nicolás de los Garza and Ciudad Apodaca, this increase seems intermittent, decreasing in a few weeks (24-31 in the case of San Nicolás de los Garza) to almost zero space fogging activities. Meanwhile, there is a permanent activity of low fogging in locations without an outbreak during most of the year.

This is the only component with a different pattern and seems not to have a significant correlation with the occurrence of the outbreak. This may be due to the weakness of space fogging as an effective component to stop transmission as some academic papers point out (Bowman et al., 2014) or due to that the fogging did not meet the necessary coverage and frequency.

## Discussion

The EWARS, developed by WHO-TDR and partner countries, has demonstrated significant prediction performace across variant settings while exceptionally allowing the use of the tool by skilled and unskilled users which can have significant operational implications (Bowman et al. 2016, Hussain Alkhateb et al 2018, WHO 2018). However, the impact of the EWARS in endemic areas remains as a crucial piece of assessment in this overall early warning and response puzzle.

Our comparative analysis using four vector control indicators from the Mexican national surveillance data, can be summarized as follows:

1. Larval control (weekly number of houses visited per week):

- In *non-outbreak* districts continuous larval control; additional activities as initial or early response
- In *outbreak districts* occasional larval control. Additional control activities came late (often as emergency response)
2. Entomological studies (number of investigations per week)

- In *non-outbreak districts* entomological studies were continuous during the entire year to monitor infestation levels. Additional activities after alarms were conducted as initial or early response
- In *outbreak districts* entomological studies were occasional and less intense. Additional control activities came late (often as emergency response)
3. Focal spraying (percentage of dengue cases with focal spraying)

- In *non-outbreak districts* interventions were continuous during the entire year suppressing the expected outbreak. Additional activities as initial or early response
- In *outbreak districts* interventions were occasional and less intense. Occasional additional activities as early or emergency response
4. Indoor residual spraying (Number of houses sprayed per week)

- In *non-outbreak districts* IRS activities were fairly continuous during the entire year helping to suppress the expected outbreak. Additional IRS as initial or early response
- In *outbreak districts* IRS activities were generally less frequent. Occasional additional activities as early or emergency response

Based on experience and historical observations from the Mexican National Surveillance and Vector Control Program, it can be argued that differences between the groups can be explained by the type and timeliness of the response to the alarms. This study applied a quantitative approach to investigate the impact of the vector control using four response indicators, comparing 11 districts grouped into ‘outbreak’ and ‘non-outbreak’ districts based on alarm signals generated by the EWARS. This investigation attempted to quantify the intensity and direction of the vector control responses before and after outbreak alarms across these districts during 2017. Two out of 11 districts eventually presented outbreaks and nine revealed alarms but no outbreaks. In overall, more intense and inclining vector controls of larval, entomological studies, spraying around cases and indoor spraying, was apparant in non-outbreak districts compared to the outbreak districts. The ‘indoor sprying’ control was however lower in terms of intensity and direction in all districts due to its nature of applications. It has to be noted that in routine vector control programs, the coverage of the whole district population is generally low because the vector control services will focus their efforts mainly on high transmission áreas (“hot spots”) where they do house-to-house visits for vector control activities. This is particularly obvious in large cities like Monterrey (with more than 1 million 200 thousand population).

Details from the segmented time series analysis showed plausible trend and pattern between and within the groups, which can further augment the assessement of the EWARS role in this vector control process. The baseline (routine) response of any of the vector control indicators appears to be crucial for defining the outbreak profile of the endemic area – since districts with poor baseline response have higher tendency for disease outbreaks. This scenario is more import in occasions where districts fail to take escalated response after a triggered early warning system alarm. Furthermore, the immediate response action at the week of declaring an alarm signal for a forthcoming outbreak plays an equally important role in redcuing the probability of having outbreak, as revealed in our results. This can assist ditrict managers to decide on the continuenity and mechanism of scaled response during the later period post-alarm signal. For instance, in the two outbreak districts, the larval control response was in the form of emergency response – when case numbers of dengue incidence had already crossed the threshold and the outbreak had started. In the non-outbreak districts, however, five out of nine had started with initial response after the first alarm followed by early response activities (when the alarm continued), two had started with initial response activities but then continued with routine larval control and two had started with early response in addition to routine activities.

While the number of outbreak district group is small (two districts), which is considered a limitation in this study, this design would still reflect a real life scenario at national and speak to the plausibility of the approach and its findings which is important for operational aspects. Some confounding variables that could not be taken into account in the analysis may further impact on the conclusion, such as the severity of the endemic districts or the periodicity of the transmission cycles, and the possible alternation of serotypes. Nevertheless, crude results from a robust segmented time series anlaysis suggest that continuous routine vector control enforced after outbreak alarm singal by additional efforts can make a difference in mitigating dengue outbreaks which can reduce further health system negative consequences.

This article neither attempts to evaluate the adherence of health district managers to the early warning and response system nor assessing the effectiveness of EWARS, which typically requires robust randomized control trials and accounts for additional factors related to resource availabilities and managerial aspects. However, it plausibly demonstrates important operational scenarios when succeeding or failing alarms signals generated by EWARS at national level. This study presents clear evidences of needs for further investigation of the effectiveness and cost-effectiveness of EWARS using gold-standard designs at national or regional levels. Subsequent evaluations could include the availability of resources at the district level, because insufficient resources can naturally affect the timeliness and coverage of actions and results.

## Availability of data and materials

Data are available from the National Center of Diseases Control and Preventive Programs (CENAPRECE) as Institutional Data Access for researchers who meet the criteria for access to confidential data.

## Authors’ contributions

DBV, AK, GST and LHA conceptualized and designed the paper which was drafted by DBV. Data curation and field implementation by DBV and GST. DBV and LHA performed the analysis. The final draft of the paper was approved by all the authors.

